# Perceptual modulation of parietal activity during urgent saccadic choices

**DOI:** 10.1101/2019.12.12.874313

**Authors:** Joshua A. Seideman, Emilio Salinas, Terrence R. Stanford

## Abstract

The lateral intraparietal cortex (LIP) contributes to visuomotor transformations for determining where to look next. However, its spatial selectivity can signify attentional priority, motor planning, perceptual discrimination, or other mechanisms. Resolving how this LIP signal influences a perceptually guided choice requires knowing exactly when such signal arises and when the perceptual evaluation informs behavior. To achieve this, we recorded single-neuron activity while monkeys performed an urgent choice task for which the perceptual evaluation’s progress can be tracked millisecond by millisecond. The evoked presaccadic responses were strong, exhibited modest motor preference, and were only weakly modulated by sensory evidence. This modulation was remarkable, though, in that its time course preceded and paralleled that of behavioral performance (choice accuracy), and it closely resembled the statistical definition of confidence. The results indicate that, as the choice process unfolds, LIP dynamically combines attentional, motor, and perceptual signals, the former being much stronger than the latter.

---

Accurate guidance of goal-directed behaviors based on incoming information from the external environment requires reliable, dynamic communication between sensory, cognitive and motor systems, and neurons in the posterior parietal cortex have long been posited to play a role in this process (Snyder et al., 2000; Gold and Shadlen, 2007; Gottlieb, 2007; Bisley and Goldberg, 2010; Hanks and Summerfield, 2017; Huk et al., 2017; Freedman and Ibos, 2018). In particular, receiving extensive visual inputs and in turn projecting to saccade production centers, the lateral intraparietal area (LIP) is aptly situated to help mediate the visuomotor transformations required for perceptually-guided saccadic choices (Petrides and Pandya, 1984; Andersen et al., 1985; Andersen et al., 1990a, 1990b; Schall et al., 1995; Paré and Wurtz, 1997; Ungerleider et al., 2007). Physiological evidence supports this hypothesis. LIP responses span a continuum from visual-to motor-related (Gnadt and Andersen, 1988; Andersen et al., 1990a, 1990b; Baizer et al., 1991; Barash et al., 1991a, 1991b; Paré and Wurtz, 1997), and across a variety of choice contexts, LIP activity has been demonstrated to discriminate targets from distracters in the response field (RF), or to otherwise categorize visual stimuli according to task rules (Gottlieb et al., 1998; Balan et al., 2008; Ipata et al., 2009; Ong et al., 2017; Zhou and Freedman, 2019). Whether interpreted as a signal of motor intent (Snyder et al., 2000), attentional deployment (Bisley and Goldberg, 2010), or sensory evidence accumulation (Gold and Shadlen, 2007), LIP activity that evolves in advance of a perceptually-informed saccadic choice is presumed to play an essential role in guiding it.

Although many studies have estimated the presaccadic time point at which such perceptually-based discrimination signals first emerge, as well as their underlying dynamics (Bisley and Goldberg, 2003; Buschman and Miller, 2007; Balan et al., 2008; Ganguli et al., 2008; Katsuki and Constantinidis, 2012; Nishida et al., 2013; Swaminathan et al., 2013; Ong et al., 2017; Sapountzis et al., 2018), these measures represent imprecise and incomplete characterizations of the temporal relationship between neurometric and psychometric performance. The present study pursues a related but more rigorous approach: we ask whether the perceptual modulation of LIP activity and the likelihood of making an accurate choice evolve with similar temporal profiles, a more direct and stringent test for how the former might guide the latter. Simply put, if LIP activity contributes to guiding perceptually informed choices, as has been suggested, then its perceptual modulation should both lead and directly parallel that of choice accuracy. We exploit an urgent choice task to accurately resolve and compare the evolution of these neural and behavioral metrics.

Beyond target-distracter discrimination, another way in which LIP neurons might contribute to perceptually guided behavior is by estimating the probability that a choice is correct given the sensory evidence — that is, by computing decision confidence as defined statistically (Berger, 1985; Hangya et al., 2016; Pouget et al., 2016; Sanders et al., 2016; Ott et al., 2019). Confidence, understood this way, is hypothesized to be a fundamental component of the perceptual decision-making process (Kepecs et al., 2008; Drugowitsch, 2016; Lak et al., 2017; Ott et al., 2019), and indeed, several of its analytically derived signatures have been observed in both behavioral and electrophysiological datasets (Kepecs et al., 2008; Kiani and Shadlen, 2009; Komura et al., 2013; Lak et al., 2014; Sanders et al., 2016; Lak et al., 2017; Urai et al., 2017; Seideman et al., 2018). However, the issue of timing is again critical to establishing a possible causal role, and almost all the neurophysiological data on confidence are based on measures taken after the choice process has already been initiated (Kepecs et al., 2008; Lak et al., 2014; Rutishauser et al., 2015). Thus, it is currently unknown whether, for any particular neural structure, sensory evidence informs the computation of decision confidence before, simultaneously with, or after the decision/choice (Kepecs et al., 2008; Fetsch, et al., 2014; Kiani et al., 2014; Drugowitsch, 2016; Lak et al., 2017; Ott et al., 2019).

Using a recently developed urgent choice task, we have shown in previous studies that the psychophysical discrimination of visual targets from distracters evolves according to how long the relevant sensory information has been viewed (Salinas et al., 2010; Stanford et al., 2010; Shankar et al., 2011; Costello et al., 2013; Salinas et al., 2014; Scerra et al., 2019), and that decision confidence may be computed within oculomotor circuits during task performance (Seideman et al., 2018). In both cases, the temporal dependence on perceptual processing time is revealed with millisecond precision. Here, we leverage the high temporal resolution afforded by this urgent paradigm to determine if modulation of single-neuron LIP activity is a direct neural antecedent to these key psychophysical quantities, choice accuracy and decision confidence.

## Results

### Spatial selectivity and feature-based target selection

We recorded from 59 neurons that exhibited spatially selective visual, delay period, and presaccadic activation within the LIP of two monkeys (Methods). When recorded in the context of single-target visually- (Fig. 1a) and memory-guided (Fig. 1b) delayed saccade tasks (see Methods), these neurons demonstrated visuomotor properties characteristic of LIP (Gnadt and Andersen, 1988; Barash et al., 1991a, 1991b; Paré and Wurtz, 1997). For both tasks, strong, spatially specific activation is clearly evident in the average population activity (Fig. 1a, b). This response is initially linked to stimulus onset, continues throughout the visual (Fig. 1a) and memory delay periods (Fig. 1b), and increases immediately prior to saccade onset. These features are typical of LIP neurons that project directly to saccade production centers, such as the superior colliculus (SC; Paré and Wurtz, 1997).

**Figure 1.**
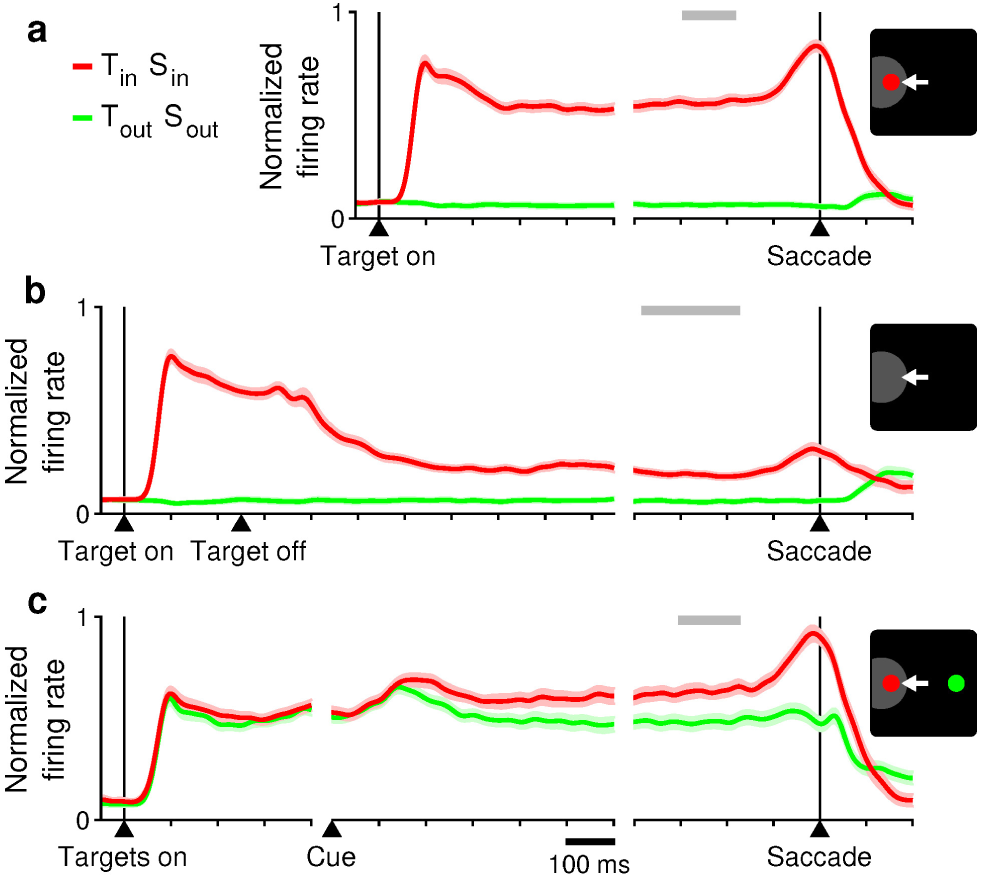
Spatial selectivity of recorded LIP activity during non-urgent tasks. Traces show normalized firing rate as a function of time for the sampled LIP population during correct saccadic choices into (red) or away from the RF (green). Shades behind traces indicate ±1 SEM across neurons. (**a**) Responses in the visually guided delayed-saccade task (*n* = 59). (**b**) Responses in the memory-guided saccade task (*n* = 59). (**c**) Responses in the delayed color discrimination task (*n* = 45). Icons depict target stimulus (red circle), distracter stimulus (green circle), and saccade (white arrow) relative to RF location (gray spot) at saccade onset. Gray bars indicate time intervals when go signals were given (90% confidence intervals). Continuous firing rates were obtained by convolving recorded single-neuron spike trains with a gaussian function (σ = 15 ms), averaging across trials, dividing by the maximum value for each neuron, and averaging across neurons. T_in_, target in; T_out_, target out; S_in_, saccade in; S_out_, saccade out.

Also consistent with prior reports (Gottlieb et al., 1998; Balan et al., 2008; Bisley and Goldberg, 2010; Ibos and Freedman, 2017), the neurons in our sample discriminated a visual target from a distracter in the RF during performance of a non-urgent perceptual discrimination task (Fig. 1c). In this ‘easy’ choice task, a red or green fixation point is followed by two gray stimuli (potential targets) and, after a delay, a color change (Cue) reveals which stimulus is the target (match to fixation point color) and which is the distracter (non-match) (see Methods for details). Neuronal activity in the easy choice task becomes selective for location only after target and distracter are revealed by the color cue (Fig. 1c), indicating that such spatial specificity is informed by the feature-based relevance of the visual stimuli that guide the eventual saccadic choice. Evidence of modest target/distracter differentiation begins approximately 150 ms after the color cue and is fully realized at the time of saccade onset (Fig. 1c).

As is typical of non-urgent choice tasks like the one just described (Fig. 1c) and those used in many prior studies (e.g., Bennur and Gold, 2011), spatial selectivity develops gradually and grows monotonically to strongly signal target location as the saccade becomes imminent. However, using an urgent variant of the same color discrimination task, we have previously shown that a fully informed saccade can occur within 120 milliseconds of color cue presentation, and that saccadic choice errors are quite rare after just 200 milliseconds of processing time (i.e., cue viewing time; Stanford et al., 2010; Shankar et al., 2011; Scerra et al., 2019). These psychophysical findings indicate that, in non-urgent tasks, most if not all of the observed growth in target-distracter differentiation reflects something other than the temporal dynamics of the perceptual judgment itself, and hence it is impossible to parse how such differential signal specifically contributes to the veracity of the saccade choice. To address this issue, we recorded from the sample of 59 LIP neurons during performance of the compelled-saccade (CS) task, an urgent-choice paradigm that yields a psychophysical readout of an evolving perceptual judgment to which perceptual modulation of neural activity can be directly compared.

### Visual evidence informs choice behavior as a function of time

In the CS task (Fig. 2a), choice performance depends fundamentally on the amount of time available to view the color cue information prior to saccade onset, what we call the raw processing time (rPT). In each trial, the disappearance of the fixation stimulus at the center of the display (the go signal; Go) instructs the participant to choose between two potential targets in the periphery (identical gray spots) within a fixed reaction time (RT) window (425 ms). However, the visual cue that distinguishes the target from the distracter is only revealed later (Cue), after an unpredictable length of time following the go signal (Gap; 25–250 ms). To perform above chance, participants must use this cue information to locate the target (match to fixation point color) and direct the impending saccade to it in the milliseconds that remain before committing to and executing a saccadic choice. By design of the task, however, this period of time (the rPT) is intrinsically variable and not always sufficient, so performance varies between chance and asymptotic. Plotting choice accuracy as a function of rPT produces the “tachometric curve,” a psychophysical performance metric that defines — with millisecond temporal precision — how much time it takes for the relevant sensory cue to inform the choice process.

**Figure 2.**
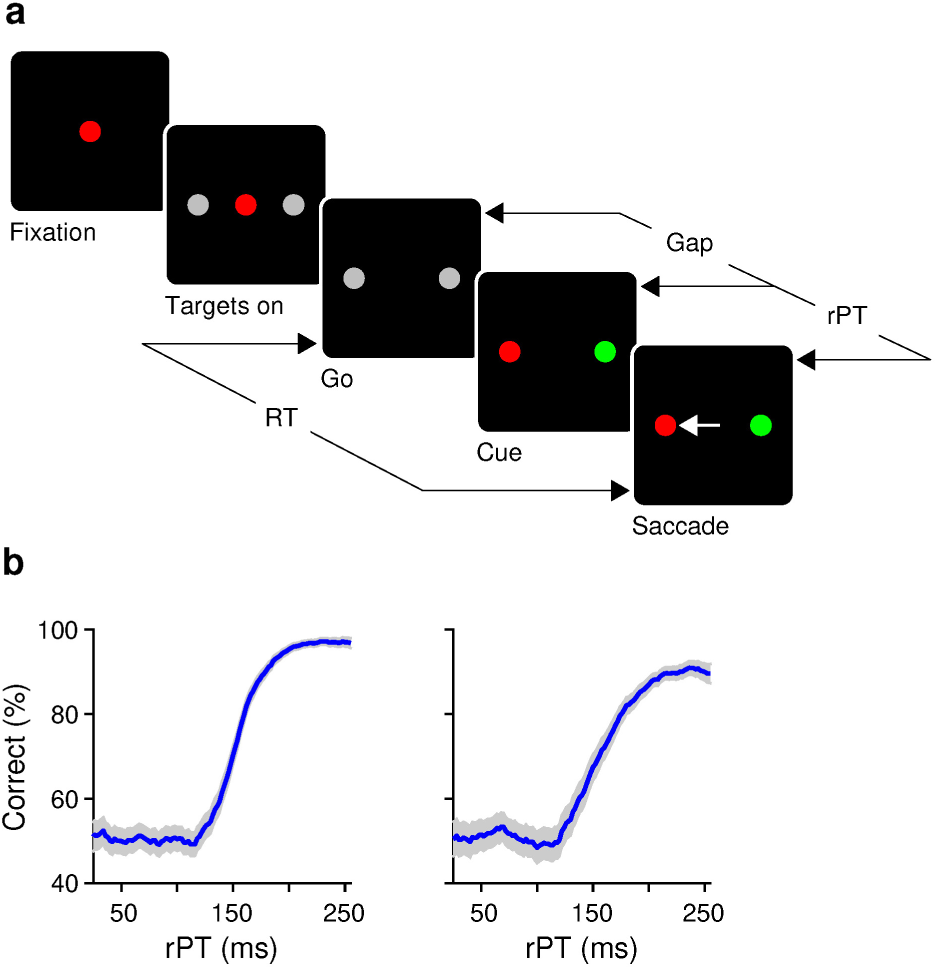
Psychophysical performance in the compelled-saccade task. (**a**) Sequence of events in the task. The imperative to respond (Go) is given *before* the color information (Cue) that identifies target (red in this example) and distracter (green), and the interval between them (Gap) is randomized (25–250 ms). Thus, the time available to view and process the color cue (rPT) varies widely from trial to trial, and so does the probability of success. (**b**) Percentage of correct responses as a function of rPT (tachometric curve) for monkey T (left) and monkey C (right). Shades represent 95% confidence intervals from binomial proportion. RT, reaction time; rPT, raw processing time.

Tachometric curves were generated for the two monkeys for the sessions in which the LIP sample was recorded (Fig. 2b). For both subjects, sensory evidence begins to inform perceptual choice behavior early, starting just ∼125 ms after the cue, and proceeds at a remarkable rate to asymptote at ∼200 ms. If, as suggested by previous work, LIP activity contributes to the generation of a visually-informed saccadic choice, then the spatial selectivity of LIP neuronal activity should evolve with a time course that directly parallels these rapid and dynamic processing-time-dependent changes in urgent perceptual performance.

### Dynamic changes in presaccadic differentiation track choice accuracy

In the CS task, LIP activity was again spatially selective (Fig. 3a, b): it was stronger for correct saccades into the RF (T_in_ choices; red traces) than for correct saccades diametrically away from the RF (T_out_ choices; green traces). Spatial differentiation was evident in the average population activity (Fig. 3b) and, to varying degrees, for individual cells (Fig. 3a). However, it was considerably weaker than that observed in the non-urgent variant of the task (Fig. 1c). To determine how much of this differentiation was informed by the color cue, we parsed trials into long- and short-rPT bins based on the tachometric curve (Fig. 3c, shaded regions) and compared LIP responses for the corresponding groups; that is, for choices that were (right panels, asymptotic performance) or were not (left panels, chance performance) guided by the cue information. The magnitude of this visuomotor signal was slightly larger for perceptually-informed choices than for guesses (Fig. 3d), as quantified via an ROC score (which compared spikes counted in the 50 ms window prior to correct T_in_ versus correct T_out_ choices; Methods). These effects, albeit modest in size, are evident upon examination of the population firing rate traces aligned to either saccade onset (Fig. 3a, b) or cue presentation (Supplementary Fig. 1). Therefore, with more time to perceptually evaluate the incoming sensory cue information, the LIP modulation prior to saccade onset becomes slightly stronger, consistent with the increased likelihood that the impending choice will be perceptually guided upon execution.

**Figure 3.**
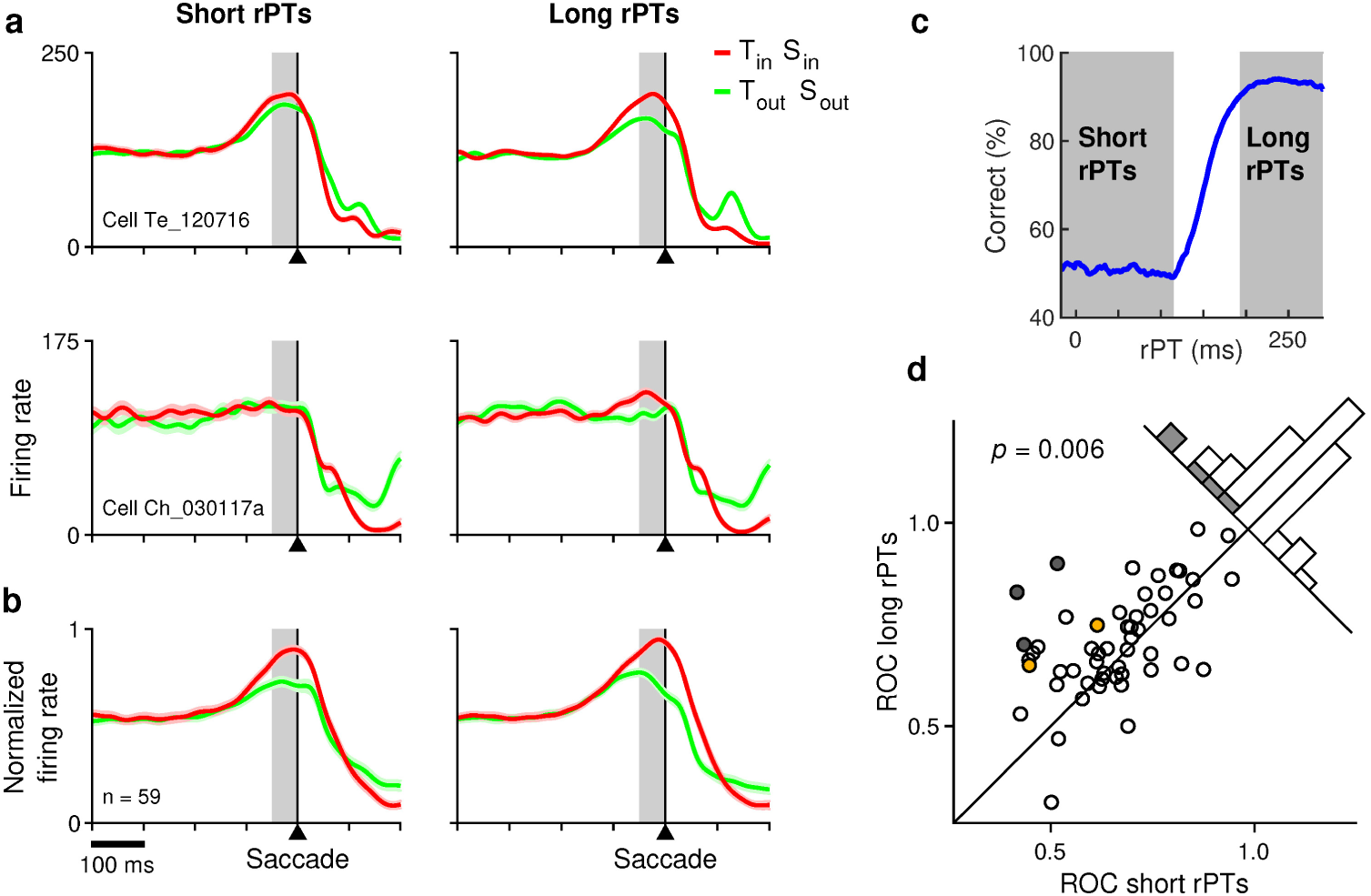
Spatial discrimination of LIP activity during urgent saccadic choices. (**a**) Responses (mean ± SEM) of two single neurons (labelled rows) recorded in area LIP. Traces show firing rate as a function of time for correct T_in_ choices (target in, saccade in; red trace) and correct T_out_ choices (target out, saccade out; green trace) made at short (left) and long (right) rPTs. (**b**) Normalized responses from the sampled population of 59 LIP neurons. Same conventions as in **a**. (**c**) Percentage of correct responses as a function of rPT (tachometric curve) for all recording sessions. Shaded regions represent short- and long-rPT groups. Note that short-rPT choices are at chance performance (uninformed), whereas long-rPT choices are highly accurate (perceptually informed). (**d**) ROC scores in long- (y-axis) versus short-rPT trials (x-axis). Each point represents data from one neuron. Significance of median difference (Wilcoxon signed rank test) is indicated. Filled symbols and gray bars mark neurons with significantly different ROC scores (*p* < 0.05, permutation test). Colored points mark the two example neurons in **a**. The ROC score quantifies the difference in LIP firing activity prior to T_in_ versus T_out_ choices (red versus green traces in **a**) based on spike counts measured before saccade onset (gray shades in **a, b** indicate 50 ms count window). Differentiation was slightly stronger prior to informed (long-rPT) than uninformed (short-rPT) choices.

To fully characterize the time course of these perceptually driven changes in LIP neuronal activity, we calculated an ROC score similar to that mentioned above (counting spikes in the same 50 ms presaccadic window), but now as a continuous function of rPT (Methods). Each point along the resulting neurometric function thus represents the degree to which T_in_ and T_out_ choices made at a given rPT can be discriminated based on the LIP neuronal responses that immediately preceded them. To directly compare the time course by which relevant sensory evidence modulates LIP activity to that by which it guides the choice, we rescaled the neurometric functions along the y-axis (Methods) and plotted them with their corresponding tachometric curves (Fig. 4).

**Figure 4.**
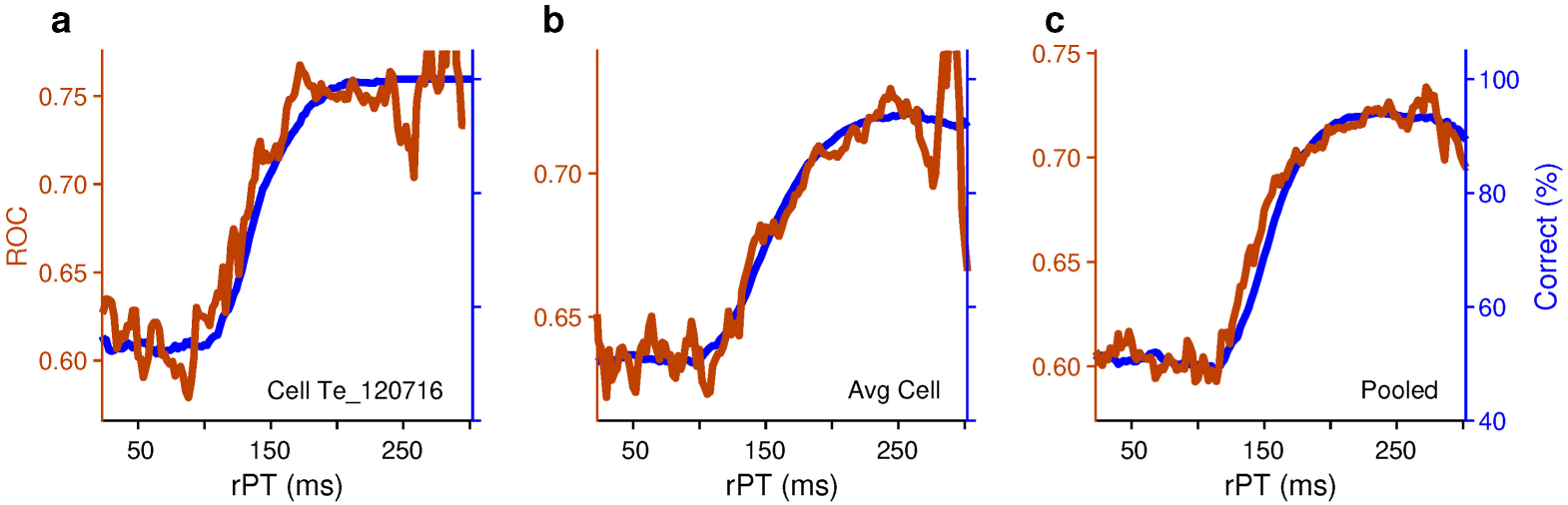
Millisecond-by-millisecond correspondence between neuronal and behavioral discrimination. Each panel plots a neurometric curve (brown trace; ROC score as a function of rPT) and a corresponding tachometric curve (blue trace; percentage of correct responses as a function of rPT) based on neural and behavioral data from the same recording session(s). ROC scores compare correct T_in_ (target in, saccade in) versus correct T_out_ trials (target out, saccade out). (**a**) Results for a single LIP neuron/recording session. (**b**) Results for the average LIP neuron/recording session. (**c**) Results for data pooled across all neurons/sessions. Data in **b** and **c** are from *n* = 59 cells.

This analysis revealed that, as a function of rPT, the LIP neuronal discriminability was, on a millisecond-by-millisecond basis, commensurate with the probability that the ensuing choices were perceptually guided upon execution. At first, discriminability remained at a relatively low and constant level, changing very little, if at all, across short rPTs (i.e., rPTs < 120 ms), similar to choice accuracy throughout the same time frame. The LIP discriminability then quickly increased to higher asymptotic levels in parallel with psychophysical performance, indicating that relevant sensory evidence modulates LIP neuronal activity at the same rapid rate that it informs the urgent choice. These results were observed in data from individual neurons (Fig. 4a), from the average neuron (Fig. 4b), and pooled across all neurons (Fig. 4c). Quantitatively, the rise times of the neurometric and tachometric curves were statistically indistinguishable (neurometric: 66 ms in [34, 140] ms; tachometric: 68 ms in [60, 75] ms; 95% confidence intervals from bootstrap for the pooled data; Methods).

### Perceptual versus motor contributions to neuronal differentiation

Next, we investigated the degree to which the observed rPT-dependent increase in LIP discriminability reflected target-distracter selectivity (a perceptual signal) or, alternatively, stronger spatial selectivity per se (a post-perceptual signal). We repeated the analyses described above but restricted to saccadic choices in a fixed direction, either toward the RF of the recorded neuron or away from it. The results indicated that, independently of the direction of the urgent saccadic choice, LIP neurons tended to fire more when the stimulus in the RF was the target rather than the distracter (Supplementary Fig. 2).

The data were clearest for identical eye movements away from the RF (Supplementary Fig. 2d, e). In that case, the LIP activity just prior to saccade onset was slightly but noticeably stronger for (incorrect) T_in_ than (correct) T_out_ choices (blue vs. green traces). Consistent with it being cue-driven, this modulation was observed only for choices made at long rPTs when perceptual information had the potential to influence presaccadic activity (Supplementary Fig. 2d, e, right panels); prior to guesses, at short rPTs, the same neurons failed to discriminate target from distracter (Supplementary Fig. 2d, e, left panels). Comparison of ROC scores computed separately for short- and long-rPT trials confirmed the robustness of this result quantitatively (Supplementary Fig. 2f). In contrast, the data for identical eye movements into the RF did not reveal a significant effect (Supplementary Fig. 2a-c). Nevertheless, the observed trend was consistent: at long rPTs the evoked presaccadic activity tended to be slightly higher for correct choices to the target than for incorrect choices to the distracter (red vs. cyan traces; Supplementary Fig. 2a, b, right panels).

The small magnitude of the cue-driven LIP differentiation is not surprising, because the selectivity of LIP neurons during urgent choices is modest to begin with (Fig. 3), and part of it must be decidedly spatial (Ipata et al., 2006, 2009). Such a weak signal is also expected to be even less detectable when the motor contribution to the evoked response is stronger — which is when the saccade is made into the RF (Kiani et al., 2008; Ipata et al., 2009). Regardless of its size, however, it is still interesting to consider whether the temporal dynamic of this cue-driven modulation is congruent with that of the behavioral choice.

To investigate this, we first selected subpopulations of neurons in order to maximize the cue-driven differential signal (Methods). Then, we once again computed ROC scores as functions of rPT, this time for saccadic choices made in a given direction (either into or away from the RF). As before, we plotted the neurometric and corresponding tachometric functions together — directly comparing neuronal changes over time in target-distracter discriminability to overt changes over time in perceptual discriminability (Fig. 5). We found that, as functions of rPT, these purely cue-driven LIP discrimination signals were also commensurate with the probability that the ensuing choices were correct. This was the case for single neurons (Fig. 5a, d) and population averages (Fig. 5b, c, e, f). As functions of rPT, the LIP signal and the monkeys’ choice accuracy increased with statistically identical rise times, both for saccades into the RF (Fig. 5c; neurometric: 32 ms in [10, 118]; tachometric: 52 ms in [44, 66]; 95% confidence intervals from bootstrap for the pooled data) and away from the RF (Fig. 5f; neurometric: 71 ms in [18, 165]; tachometric: 62 ms in [53, 72]). Taken together, the results indicate that, although small, the perceptually-driven changes in LIP activity precede and evolve in parallel with concomitant changes in urgent perceptual performance.

**Figure 5.**
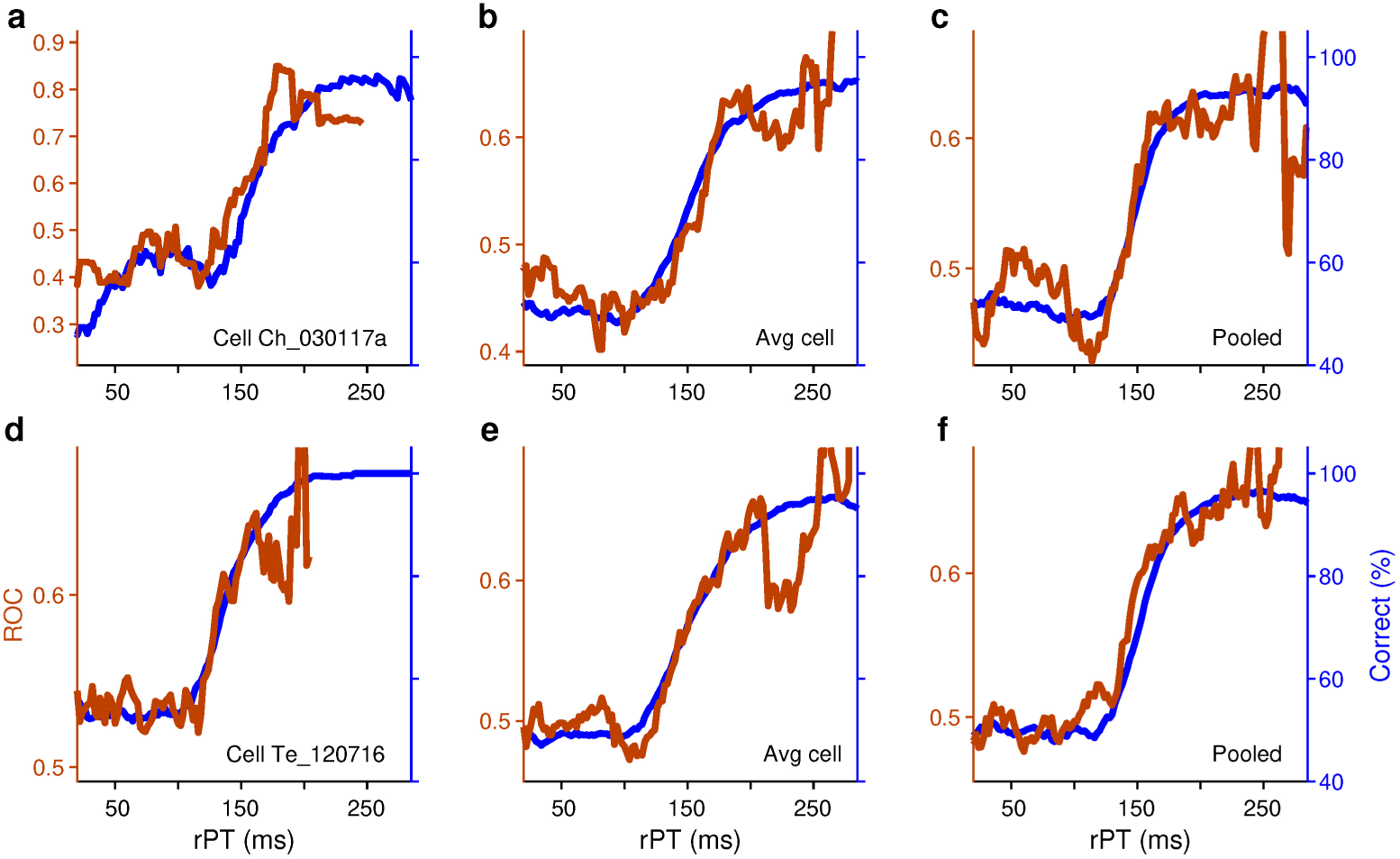
Parietal target-selection dynamics are independent from movement direction. Each panel plots a neurometric (brown trace) and a tachometric curve (blue trace), as in Fig. 4, but for saccadic choices in a given direction. (**a**–**c**) Results for trials that resulted in saccades into the RF, collected from *n* = 14 neurons/sessions (Methods). ROC scores compare correct T_in_ (target in, saccade in) versus incorrect T_out_ trials (target out, saccade in). Tachometric curves are based on all trials from the corresponding recording sessions. Results are shown for a single LIP neuron (**a**), the average neuron (**b**), and pooled across neurons (**c**). (**d**–**f**) Results for trials that resulted in saccades away from the RF, collected from *n* = 21 neurons/sessions. ROC scores compare incorrect T_in_ (target in, saccade out) versus correct T_out_ trials (target out, saccade out). Same format as in **a**–**c**. The neural activity in LIP evolves to discriminate target from distracter in parallel with urgent choice accuracy.

### Oculomotor correlates of statistical decision confidence

Recent work from our laboratory (Seideman et al., 2018) suggests that, in the CS task, neurons within oculomotor structures compute the probability that an impending choice will be correct given the sensory evidence — i.e., they compute decision confidence as defined statistically. We therefore investigated whether LIP neurons might reflect or participate in this computation. For this analysis, the same subpopulations used in Fig. 5 were considered.

We found that LIP neuronal activity recorded prior to urgent choice onset exhibits three analytically proven signatures of confidence (Fig. 6; Hangya et al., 2016). For saccades made into the RF, higher firing activity corresponded to higher confidence, and the three signatures were as follows. First, the evoked LIP responses correlated positively (and strongly) with choice accuracy (Fig. 6a, g; *r* = 0.98, *p* = 0.002 for the pooled data). In other words, LIP activity predicted the accuracy of the perceptual choices that ensued. Second, on average, the spike counts measured before correct saccades increased as functions of rPT, whereas those measured before incorrect saccades decreased (Fig. 6b, h). And third, the tachometric curve shifted when conditioned on LIP activity, such that psychophysical performance was enhanced when the evoked spike counts were high compared to when they were low (Fig. 6c, i). In contrast, for saccades made away from the RF, *lower* firing activity corresponded to higher confidence (or equivalently, higher activity corresponded to higher decision uncertainty, which is the opposite of confidence; Kepecs et al., 2008; Kiani and Shadlen, 2009; Urai et al., 2017). The representation of confidence in this case was perfectly complementary to that for saccades into the RF, as the opposite modulation patterns were found for the three signatures (Fig. 6d-f, j-l; *r* = –0.97, *p* = 0.003 for the pooled data in panel j).

**Figure 6.**
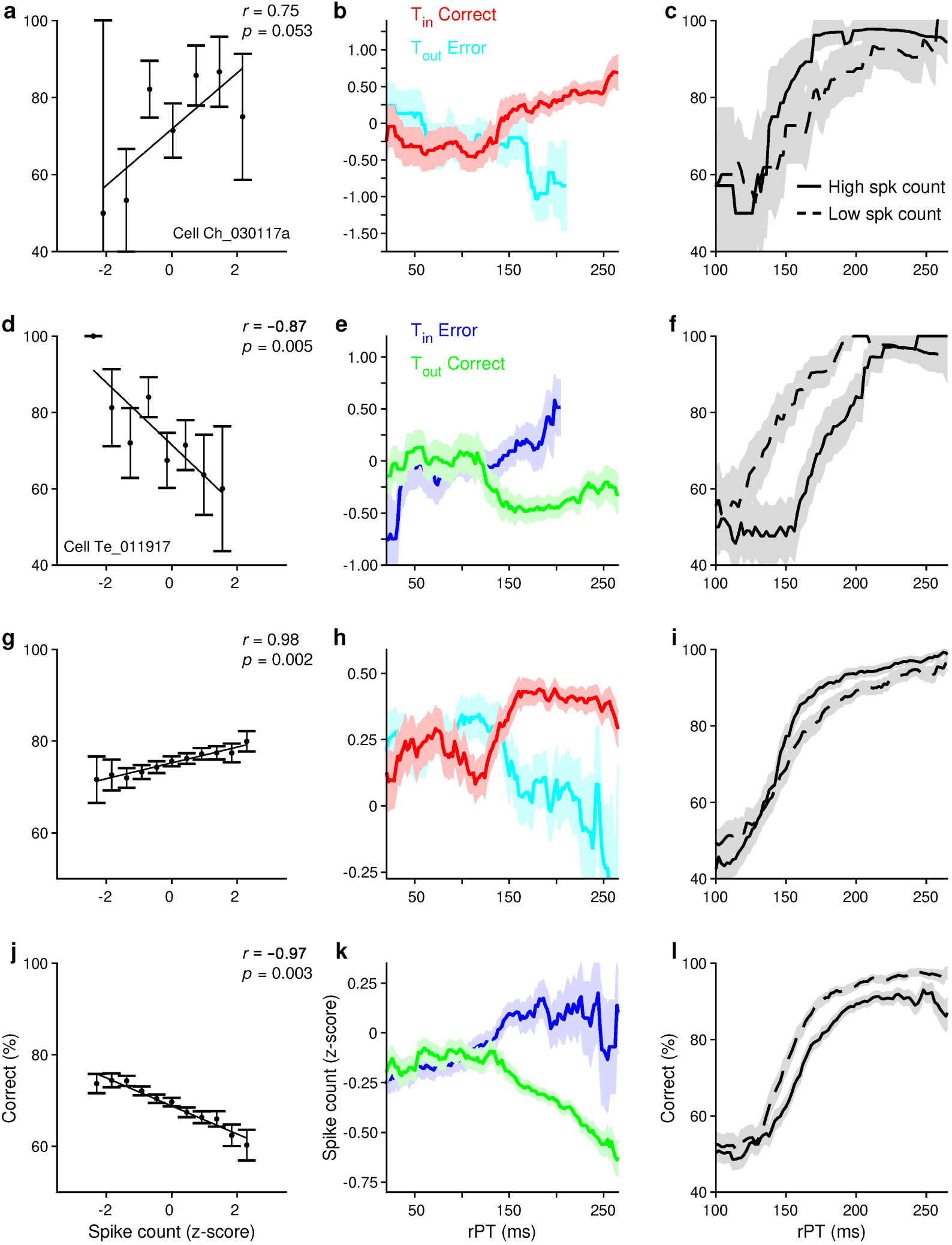
Oculomotor correlates of decision confidence. (**a**–**c**) Activity from a single neuron exhibits characteristic features of confidence, as defined statistically. (**a**) Percentage of correct responses (± SEM) as a function of spike count measured prior to choices into the RF. Pearson correlation (*r*) between values on the x and y axes is indicated, along with *p*-value (Methods). (**b**) Mean spike count (± SEM) as a function of rPT for correct (red; target in, saccade in) and incorrect choices (cyan; target out, saccade in) executed into the RF. (**c**) Percentage of correct responses (± SEM) as a function of rPT (tachometric curves) for choices into the RF with spike counts that fell above (solid line) versus below (dashed line) the median. (**d**–**f**) same format as in (**a**–**c**), but for correct (target out, saccade out; green traces in **e**) and incorrect choices (target in, saccade out; blue traces in **e**) executed away from the RF. Data are from another single neuron. (**g**–**l**) Population results based on the same two groups of cells analyzed in Supplementary Fig. 2 and Fig. 5. (**g**–**i**) Same format as in (**a**–**c**) but for data pooled across 14 neurons. (**j**–**l**) Same format as in (**d**–**f**) but for data pooled across 21 neurons.

These results indicate that the LIP activity evoked before saccade onset contains a representation of statistical decision confidence that is congruent with LIP’s spatial selectivity. Moreover, given that confidence reports were in no way explicitly solicited by the task, and that the confidence signal followed the same time course as performance (note correspondence between Figs. 5c, f and 6h, k), it appears that this representation of confidence arises naturally as part of the choice process itself as it develops.

## Discussion

We investigated if, how, and when incoming perceptual information influences LIP neuronal activity under conditions in which the ability to accurately guide a saccadic choice depends critically on sensory cue processing time, much like eye movements made under natural time pressure. We found that, in the urgent CS task, perceptually-driven changes in LIP activity preceded and evolved in parallel with concomitant changes in the perceptual discriminability of the choice alternatives. In addition, this LIP response exhibited multiple features that are characteristic of the statistical definition of confidence (i.e., the probability that a hypothesis is correct given the evidence), even though confidence estimates were not explicitly required by the urgent task. Although the results demonstrated a direct, millisecond-by-millisecond correspondence between LIP activity and the temporal evolution of a perceptual judgment, they also revealed that the magnitude of the LIP modulation specifically attributable to perceptual information is quite meager. Contrasted with the robust choice-related differentiation observed under more relaxed response-time constraints, this finding suggests that most of the spatial differentiation observed in “non-urgent” tasks is driven by factors unrelated to the perceptual judgment itself.

### Presaccadic perceptual modulation of LIP activity parallels choice accuracy

Previous studies have employed a variety of tasks in efforts to either characterize the temporal dynamics of perceptually-based modulation of LIP activity (Bisley and Goldberg, 2003; Buschman and Miller, 2007; Balan et al., 2008; Ganguli et al., 2008; Katsuki and Constantinidis, 2012; Nishida et al., 2013; Swaminathan et al., 2013; Ong et al., 2017; Sapountzis et al., 2018) or determine the amount of time needed for relevant cue information to guide a choice (Afacan-Seref et al., 2018; also see Gold and Shadlen, 2000; Katnani and Gandhi, 2013). As detailed in our prior studies (Salinas et al., 2010; Stanford et al., 2010; Shankar et al., 2011; Costello et al., 2013; Salinas et al., 2014; Seideman et al., 2018; Scerra et al., 2019), the CS task used here has two distinct advantages for relating visuomotor activity to behavioral performance. First, the resulting tachometric curve is an overt behavioral expression of a developing perceptual judgment, with each point reflecting a state to which neural activity may be directly compared. As such, any parallel between the tachometric curve and the magnitude of neural differentiation over time represents a millisecond-precision test of the veracity with which neural activity correlates with the ensuing perceptually guided choice. Second, comparison of the activity preceding informed versus uninformed saccades that are otherwise metrically identical provides an opportunity to distinguish the specific contributions of perceptual versus non-perceptual (e.g., motor) processes to the spatial selectivity that precedes the choice. Here, any difference in neural differentiation for chance (uninformed) versus asymptotic (informed) performance delineates the upper limit for the contribution of perceptual evidence (i.e., color, in this case) to target selection.

We found a very tight temporal correspondence between choice accuracy and the rPT-dependent growth in LIP spatial differentiation for the activity epoch immediately preceding the saccade. Accordingly, our findings indicate that LIP neurons are modulated by relevant sensory cues in a time frame that is consistent with its proposed role in guiding perceptual choice behavior. This need not have been the case. In a previous study of the frontal eye field (FEF), we reported on a class of neuron that strongly selected salient targets under non-urgent conditions, but did so less vigorously when the discrimination was made urgent (Scerra et al., 2019); and furthermore, when the discrimination was between a distracter and an equally salient target, as in the current experiment, those same neurons failed to select the target at all for informed choices (Costello et al., 2013; Scerra et al., 2019). Thus, robust spatial selection in a non-urgent task with one stimulus configuration (e.g., Fig. 1) does not necessarily predict the same for a different configuration, or even for its urgent counterpart. It is therefore conceivable that the rPT-dependent modulation in LIP activity would have failed to manifest at all or, given the stringent temporal constraints of the CS-task, perhaps lagged that of choice accuracy. Although the observed temporal correspondence between LIP modulation and the tachometric curve is consistent with the idea that sensory evidence informs choice behavior by way of LIP, we note that such correlative findings cannot rule out the possibility that LIP activity represents a copy of the decision-relevant information, but within a circuit separate from that necessary to guide the choice (Katz et al., 2016; Pisupati et al., 2016; Subramanian et al., 2019).

### Non-perceptual factors predominate in accounting for LIP spatial selectivity

Traditionally, spatial selectivity in LIP has been interpreted as deployment of internally-derived spatial guidance mechanisms, such as those relating to motor intention or spatial attention (Goldberg et al., 1990; Colby et al., 1996; Snyder et al., 1997, 2000). Many subsequent studies have elaborated on these putative roles in describing LIP signals related to urgency (Churchland et al., 2008; Drugowitsch et al., 2012; Hanks et al., 2014) or attentional priority (Gottlieb et al., 1998; Kusunoki et al., 2000; Bisley and Goldberg, 2003; Balan and Gottlieb, 2006; Buschman and Miller, 2007; Bisley and Goldberg, 2010). The present findings are consistent with either account and, in this regard, the substantial difference in LIP spatial selectivity associated with otherwise identical urgent and non-urgent informed choices is instructive.

In the non-urgent condition, presaccadic activation for saccades into the RF far exceeded that for saccades away from the RF, an unambiguous selection of the target that developed over hundreds of milliseconds and which peaked just prior to saccade execution. In contrast, activity in the urgent condition was characterized by spatial conflict that persisted to within 100 milliseconds of saccade onset and which was resolved to a much lesser extent at the time of saccade execution. As discussed in prior studies, such conflict is strongly promoted by urgent paradigms (Chapman et al., 2010; Stanford et al., 2010; Costello et al., 2013; Gallivan et al., 2015; Scerra et al., 2019) and likely reflects the simultaneous planning of multiple movement options (Cisek and Kalaska, 2005, 2010; Scherberger and Andersen, 2007; Thura and Cisek, 2014) and/or the division of spatial attention across equally salient potential target locations (Awh and Pashler, 2000; Bisley and Goldberg, 2003; Godijn and Theeuwes, 2003; McMains and Somers, 2004). Assuming the latter, the timely resolution of this conflict could reflect the processing-time-dependent allocation of attentional priority away from the distracter and toward the target location.

Though strongly correlated with performance, target/distracter differentiation for informed choices was enhanced little beyond that for uninformed guesses. This relatively weak influence of perceptual information on LIP visuomotor activity seemingly conflicts with data from many previous reports utilizing delayed-response or reaction time versions of perceptual choice tasks (Roitman and Shadlen, 2002; Balan et al., 2008; Kiani and Shadlen, 2009; Bennur and Gold, 2011; Katsuki and Constantinidis, 2012; Meister et al., 2013; Nishida et al., 2013; Swaminathan et al., 2013; Hanks et al., 2014; Zhou and Freedman, 2019; although see Buschman and Miller, 2007). However, it is important to note that non-urgent tasks pose a problem for determining when and how much perceptual and non-perceptual factors contribute to the growth of activity in favor of a saccadic goal. For example, as noted for the delayed two-choice task (Fig.1c), activity profiles for target and distracter evolved for the entirety of the extended period between cue delivery and saccade onset — but based on these data alone one cannot know the degree to which differentiation reflects the consideration of perceptual evidence, spatial attention, and/or motor planning at any given time point during its progression. That said, we know from CS task performance that under urgent conditions the same perceptual judgment can be accomplished within 120–200 milliseconds. Assuming that this perceptual process unfolds similarly in the non-urgent case (i.e., is completed within 200 milliseconds), we might reasonably conclude that the large majority of the differentiating period leading up to the saccade reflects post-decision processes of intention or attention. Whether a correlate of intention or attention, the observation that target selection immediately preceding equally informed choices was considerably greater for non-urgent than for urgent choices suggests that some modulation of LIP activity is superfluous, in that it is neither necessary for perceptual choice guidance (Katz et al., 2016; Pisupati et al., 2016; Subramanian et al., 2019) nor saccade execution (Mooshagian and Snyder, 2018).

### Confidence as inherent to the decision-making process

Decision confidence represents a forecast about a choice, namely, the probability that said choice is correct given the (perceptual) evidence that it is based on. Three general interrelations between accuracy, evidence, and confidence have been identified by mathematical arguments: (1) confidence is proportional to choice accuracy, (2) for correct choices, average confidence increases with increasing evidence discriminability (i.e., strength), whereas for incorrect choices it decreases, and (3) confidence predicts outcome beyond evidence discriminability alone; that is, distinct psychometric curves are generated when trials are conditioned on confidence (Hangya et al., 2016; Pouget et al., 2016; Sanders et al., 2016; Lak et al., 2017; Ott et al., 2019). An important caveat has been noted, however. For these interrelations to qualify as signatures of the computation of confidence, certain statistical conditions must be satisfied. Specifically, the counterintuitive behavior of confidence during error trials (i.e., lower confidence for stronger evidence) is expected only when there is no overlap between stimulus distributions, so the mapping between evidence and correct choice is unambiguous (Adler and Ma, 2018), and the evidence guiding the choices does not contain independent trial-by-trial information about its discriminability (Rausch and Zehetleitner, 2018). Critically, in the CS task target and distracter are perfectly distinct, and both performance and target-distracter discriminability are determined by the same variable, processing time (rPT), so there is no further source of information about stimulus discriminability across trials. The above signatures should indeed be diagnostic of confidence.

In a previous study (Seideman et al., 2018), we found that the peak velocity of saccades in the CS task behaved very much as a confidence signal: it increased monotonically with choice accuracy, varied as a function of rPT in opposite directions for correct and incorrect choices, and produced shifted tachometric curves upon conditioning. Those behavioral manifestations of confidence were replicated by a race-to-threshold model formulated earlier (Stanford et al., 2010; Shankar et al., 2011), which also replicated frontal eye field activity recorded during the task (Costello et al., 2013). Thus, the findings of Seideman et al. (2018) strongly suggested that decision confidence is computed within oculomotor circuitry during performance of the CS task. Indeed, here we provide direct empirical support for this hypothesis, as the LIP activity recorded prior to urgent-choice onset exhibited the same qualitative signatures of confidence (compare Fig. 6 to Fig. 8 of Seideman et al., 2018). Our results are in line with an earlier study that demonstrated a correlation between LIP activity and task-solicited, post-decisional reports of confidence (Kiani and Shadlen, 2009) — except that in the CS task confidence reports were never solicited. Although such confidence information may be functionally significant (for instance, for task learning; Pouget et al., 2016), it is also possible that it arises naturally within motor selection circuits simply because perceptual evidence (input) and choice signals (output) coexist there. Either way, our results suggest that the neural computation of statistical decision confidence within LIP is a natural antecedent to perceptually-guided saccadic choices.

## Methods

### Experimental model and subject details

All experimental procedures were conducted in accordance with NIH guidelines, USDA regulations, and the policies set forth by the Institutional Animal Care and Use Committee (IACUC) of Wake Forest School of Medicine. Two adult male rhesus monkeys (*Macaca mulatta*) weighing between 8.5 and 11 kg were the subjects in this experiment. Both animals were pair-housed in Allentown quad format cages that met all regulatory requirements. No surgical procedures had been performed on either of the animals prior to the start of this investigation. For the current study, an MRI-compatible post (Crist Instruments, MD, USA; titanium for Monkey T, polyetheretherketone for Monkey C) was implanted on the skull of each animal while under general anesthesia. The post served to fix the position of the head during all experimental sessions, facilitating data acquisition and behavioral training. Following head post implantation, both subjects were trained to perform oculomotor response tasks in exchange for water reward. After reaching a criterion level of behavioral performance (> 90% accuracy for each task), craniotomies were made and recording cylinders (Crist Instruments, MD, USA) were placed over the left LIP of each monkey (stereotactic coordinates: 5 posterior, 12 lateral; Colby et al., 1996; Snyder et al., 1998) while under general anesthesia, to facilitate neural recordings. Neural recordings commenced after a 1–2 week recovery period following cylinder placement.

To ensure that each subject maintained a healthy body weight throughout the course of the study, each subject’s weight was frequently measured and compared to a pre-determined, non-experimental baseline. Food was provided ad libitum while in their home cage. To further ensure their physical and psychological well-being, subjects were provided with food treats, manipulable objects/toys, and television on a regular basis while in their home cages.

### Behavioral and neurophysiological recording systems

Eye position was monitored using an EyeLink 1000 Plus infrared tracking system (SR Research; Ottawa, Canada), operating with a sampling rate of 500 Hz. Gaze-contingent stimulus presentation and reward delivery were accomplished via a custom-designed PC-based software package (Ryklin Software). Visual stimuli were presented on a Viewpixx/3D display (Vpixx Technologies, Quebec, Canada; 1920 × 1080 screen resolution, 120 Hz refresh rate) placed 57 cm away from the subject.

Neural activity was recorded using single tungsten microelectrodes (FHC, Bowdoin, ME; 2–4 MΩ impedance at 1 kHz) driven by hydraulic microdrive (FHC). Microelectrodes were supported by a guide tube penetrating the dura. Electrical signals passing through the microelectrode were referenced to ground. A Cereplex M headstage (Blackrock Microsystems, Utah, USA) filtered (0.03 Hz–7.5 kHz), amplified, and digitized electrical signals, which were then sent to a Cereplex Direct (Blackrock Microsystems) data acquisition system. Single neurons were isolated online based on amplitude criteria and/or waveform characteristics.

### Behavioral tasks

#### Delayed visually-guided and memory-guided saccade tasks

We used two single-target saccade tasks to characterize the essential visuomotor properties of neurons within the LIP sample. For both tasks, a trial begins with presentation of a central fixation spot. Upon fixation and after a short delay, a peripheral target is presented (Target on) either within or diametrically opposed to the response field (RF) of the recorded neuron. For the delayed visually-guided saccade task, the fixation spot disappears (Go signal) after a variable delay (500–1000 ms) and the monkey is required to make a saccade to the peripheral target within 600 ms to obtain a liquid reward. For the memory-guided saccade task, the peripheral target is extinguished before the Go signal (Target off), and the monkey is required to maintain fixation throughout a subsequent delay interval (500–1000 ms). This memory retention interval concludes with offset of the fixation spot (Go signal), thus signaling the monkey to make a saccade to the remembered location of the peripheral target within 600 ms to obtain a liquid reward.

#### Delayed choice task

The delayed choice task is a two-alternative task which requires the monkey to discriminate a target from a distracter stimulus on the basis of color. Each trial begins with presentation of a central fixation spot whose color (red or green) defines the identity of the eventual target. Upon fixation and after a short delay (300–800 ms), two gray stimuli (potential targets) are presented (Targets on), one in the RF and one diametrically opposed. After a delay (250–750 ms), one of the gray stimuli changes to red and the other to green (Cue). After an additional delay period (500–1000 ms), the fixation spot is extinguished and the monkey is required to make a saccade to the stimulus that matches the color of the prior fixation spot within 600 ms to obtain a liquid reward. Colors and locations for target and distracter are randomly assigned in each trial.

#### Compelled-saccade (CS) task

As deployed here, the CS task requires the same red/green color discrimination as in the easier delayed choice task. The key distinction is that the CS task mandates an urgent decision/choice by limiting the amount of time available to process the color information before committing to a saccade. Each trial of the compelled-saccade task (Fig. 2a) begins with the presentation of a spot at the center of the display, the color of which (red or green) defines the color of the eventual correct target. Once the monkey fixates on the central spot (Fixation; 300–800 ms), two gray stimuli (potential targets) are presented in the periphery (Targets on), one in the RF of the recorded LIP neuron and one diametrically opposed. Then, after 250–750 ms, the fixation spot disappears (go signal; Go), instructing the monkey to make a choice to one of the potential target stimuli. The go signal urges the subject to respond as quickly as possible because, if a saccade is not initiated within a fixed time window (approximately 425 ms), the trial times out and no reward is obtained. At this point in the trial, however, no information is available to guide the choice above chance performance; that is, one of the remaining stimuli is the correct target, yet there is no way to identify it (50% of the responses made during this task epoch are randomly classified as correct). Rather, if time permits, the sensory cue necessary for informing the choice is revealed later (one gray spot turns red and the other green; Cue), after an unpredictable length of time following the go signal (Gap; 25– 250 ms). Subjects are tasked with looking to the peripheral choice alternative whose color matches that of the initial fixation spot (Saccade). A correct saccadic response is rewarded with a drop of water. The location and color of the correct target varies randomly from trial to trial. It is important to note that, in the CS task, there is neither an explicit requirement nor an incentive to report or estimate the confidence associated with a decision/choice.

The reaction time (RT) is measured as the time elapsed between the disappearance of the fixation spot and the initiation of the saccade. On each trial, the rPT is measured as the duration of time between cue onset and saccade onset (rPT = RT – gap). For example, a choice made at 75 ms rPT means that the saccade was initiated 75 ms after cue onset. Task difficulty is controlled by manipulating the gap length, which has a largely complementary relationship with the rPT (Salinas et al., 2010; Stanford et al., 2010; Shankar et al., 2011; Costello et al., 2013; Seideman et al., 2018; Scerra et al., 2019). That is, trials with short gaps are more likely to result in long-rPT responses, which are typically correct, and trials with long gaps are more likely to result in short-rPT responses, which are typically at chance. The gap duration (25-250 ms) varies randomly from trial to trial. Gap values were chosen to yield rPTs covering the full range between guesses and fully informed choices.

### Analysis of behavioral data

All data analyses were performed in Matlab (The MathWorks, Natick MA). Saccade onset was determined as the time point at which eye velocity exceeded 25°/s. For multiple neural analyses, trials were parsed into short and long rPT time bins based on the tachometric curve (e.g., Fig. 3, and Supplementary Figs. 1, 2). To define these two rPT intervals, tachometric curves were first fit with a piece-wise-linear version of a sigmoid function to estimate the time points at which the tachometric curve started (x_1_) and finished (x_2_) rising (Seideman et al., 2018). For the neural analysis of correct T_in_ versus correct T_out_ trials, short and long rPT bins were defined as illustrated in Fig. 3c: for the short bin, rPT < x_1_ (chance performance), and for the long bin, rPT > x_2_ (asymptotic performance). For the neural analyses of T_in_ versus T_out_ trials with matching saccade directions, short and long rPT bins were defined based on the midpoint of the tachometric curve, x_m_ = (x_2_+x_1_)/2, in order to include more trials within each condition. In this case, rPT < x_m_ for the short bin and rPT > x_m_ for the long. For the scatter plot shown in Fig. 3d, short and long rPT bins were determined using the aforementioned procedure, but applied separately to each experimental session analyzed. Neurometric functions for individual neurons, which were constructed from subsets of trials from a recording session (e.g., correct T_in_ and correct T_out_ trials; see below), were compared to the full tachometric curve based on all the completed trials from that session. A similar procedure applied to the data pooled across multiple sessions.

### Characterization of neural activity

RF location was determined from activity levels measured during performance of the CS task around the time of saccade onset. Continuous firing rate traces (or spike density functions) for each neuron (as in Fig. 3a) were generated by aligning the spike trains to relevant task events (e.g., cue onset, saccade onset), convolving them with a gaussian kernel (σ = 15 ms), and averaging across trials. Normalized population traces (as in Figs. 1, 3b) were generated by dividing each cell’s response curve by its maximum firing rate value and then averaging across cells. For each cell, the same maximum firing rate value (calculated from activity recorded during performance of the CS task) was used to normalize the population traces for all behavioral tasks.

All neurons included in the current study (n = 59) were significantly activated both in response to visual stimuli presented in their RF (window: 20:150 ms, aligned on targets on) as well as prior to saccades executed into their RF (window: –110:–10 ms, aligned on saccade), relative to respective baseline measures (visual baseline window: –150:0 ms, aligned on targets on; motor-related baseline window: –50:50 ms, aligned on go signal) during performance of the CS task. In addition, all neurons included exhibited significant delay period activity during performance of visually- and/or memory-guided saccade tasks (window: from 300 ms after target onset/offset until end of delay period; baseline window: –150:0 ms, aligned on target onset). A few additional neurons that were also recorded had no significant visual (*n* = 1), delay (*n* = 1), or presaccadic activation (*n* = 8), and were excluded from the studied sample. Significance (*p* < 0.01) was estimated via permutation test (20,000 iterations; Siegel and Castellan, 1988). These physiological response properties (i.e., visual, delay period, and presaccadic activation) are characteristic of LIP neurons that project directly to saccade production centers (i.e., the SC; Paré and Wurtz, 1997).

### ROC analyses and neurometric curves

In the current study, the area under the receiver operating characteristic (ROC) curve (Green and Swets, 1966; Fawcett, 2006) was used to quantify the degree to which LIP neurons were differentially activated across two conditions, T_in_ and T_out_ choices. This quantity, which we refer to as the ROC score, corresponds to the accuracy with which an ideal observer can classify data samples from two distributions (of responses in T_in_ and T_out_ trials, in this case). Values of 0.5 correspond to distributions that are indistinguishable (chance performance, full overlap), whereas values of 0 or 1 correspond to fully distinguishable distributions (perfect performance, no overlap). All ROC scores were computed using spike counts measured prior to choice onset (–50:0 ms, relative to saccade onset), z-scored within each recording session analyzed, and sorted according to trial outcome. Single-cell neurometric functions (Figs. 4a, 5a, 5d) were obtained by calculating an ROC score as a function of rPT (bin width = 85 ms, step size = 2 ms) for each recorded neuron. Cell-averaged neurometric functions (Figs. 4b, 5b, 5e) were obtained by then averaging across neurons. To compute pooled neurometric functions (Figs. 4c, 5c, 5f), the z-scored spike counts were first pooled across all sessions and then an ROC score was calculated as a function of rPT (bin width = 50 ms, step size = 2 ms). The bin width and step size used to compute each neurometric function were always equal to those used to compute the corresponding tachometric curve it was directly compared with. It is important to note that although the ROC scores that make up each neurometric curve vary with rPT (which is calculated based on the timing of the choice), they are always based on spike counts measured just prior to each choice.

To visually compare the time course of perceptually-driven LIP modulation to that of psychophysical performance (Figs. 4, 5), we shifted and rescaled the y-axis of each neurometric function to best match its corresponding tachometric curve. To do this, first we varied the baseline (i.e., vertical offset), the scale of the y-axis, and the origin of the x-axis (i.e., horizontal offset) of the neurometric curve until the mean absolute difference between the two curves was minimized (Seideman et al., 2018). Then the optimal y-axis shift and scaling factor resulting from the minimization solution were applied, leaving the x-axis intact. This way, the resulting rescaled neurometric function had a similar baseline and varied along a similar range in the y direction as its corresponding tachometric curve.

To determine whether incoming sensory evidence modulated LIP activity and behavioral performance at similar rates, we estimated and compared the rise times of neurometric and tachometric functions as follows. First, each curve was fitted with a piece-wise-linear sigmoid function to estimate the time points, x_1_ and x_2_, at which it started and finished rising (as described above), and the curve rise time was defined as the difference x_2_–x_1_. Then a distribution of rise times was generated by bootstrapping (Davison and Hinkley, 2006; Hesterberg, 2015); that is, by repeatedly resampling with replacement the data from each curve (2,000 iterations), refitting, and recomputing x_2_–x_1_ each time. Finally, 95% confidence intervals (CIs) calculated from these distributions were compared.

To examine the activity evoked during direction-matched saccades, neurons were analyzed and selected as follows. On a cell-by-cell basis, trials were sorted according to rPT (short or long) and saccade direction (saccades into or away from the RF). Then, for each of the four resulting groups of trials, an ROC analysis was performed comparing z-scored spike counts for T_in_ versus T_out_ choices. For a given saccade direction, only cells with more than two trials in each condition were considered. Based on this criterion, 29 neurons with saccades into the RF and 42 with saccades away were analyzed (these two groups correspond to the points in Supplementary Fig. 2c and f, respectively). ROC scores were computed such that values greater than 0.5 always corresponded to higher activity for a target compared to a distracter in the RF. Within each saccade-direction condition, the difference between the resulting ROC values measured at long- and short-rPT bins was then used as an index of rPT-dependent, target-distracter activity modulation for each cell. For each such index, significance was computed based on a permutation test in which the “short” and “long” trial labels were shuffled (2,000 iterations; filled vs. open symbols in Fig. 3d, and Supplementary Fig. 2c, f).

Within these two groups with 29 and 42 neurons, only those cells with modulation indices greater than the median value were included in the subsequent analyses of direction-matched responses, which examined their time course (Fig. 5) and relationship to decision confidence (Fig. 6). This was to isolate a relatively strong target-distracter differential signal, and resulted in subpopulations of 14 and 21 neurons for saccade-in and saccade-away conditions, respectively. To ensure that the resulting time courses (Fig. 5) were not a trivial consequence of this selection procedure, we performed the following control analysis on each of the pooled neurometric curves (Fig. 5c, f). For each curve, the z-scored spike counts of the trials were randomly permuted, breaking any possible association between spike count and rPT, as well as spike count and choice outcome (or target location). Then, modulation indices were recomputed for each cell, and cells with modulation indices greater than the median were selected for further analysis. Based on the neurons/trials thus selected, the neurometric and tachometric curves were recomputed and a Pearson correlation coefficient (between ROC and choice-accuracy values) was calculated to quantify the similarity or overlap between them. This procedure was repeated 2,000 times with different permutations of the z-scored spike count labels to generate a null distribution of coefficients, i.e., the distribution expected from limited data sampling alone, without any true association between neural activity and choice accuracy. The Pearson’s correlation coefficient for the original pooled neurometric and tachometric curves was compared to this distribution and found to lie outside its 95% CI (*p* = 0.024 for saccades in; *p* = 0.004 for saccades away; one-tailed t-tests). This demonstrates that the tight temporal correspondence between the continuous neurometric and tachometric curves was not an artifact of the neural selection procedure, which was based on a categorical distinction in activity between short- and long-rPT trials.

### Statistical confidence analyses

For pooled datasets, the average spike count as a function of rPT was computed with a bin width of 50 ms and a step size of 2 ms for each condition (i.e., Saccade in, Target in; Saccade in, Target out; Saccade out, Target in; Saccade out, Target out). The percentage of correct responses as a function of spike count was calculated using a bin size for the spike counts equal to one sixth their range and a step size equal to one quarter the bin size. The corresponding correlation was assessed using a Pearson coefficient with significance (p-value) obtained from permutation tests (10,000 iterations). To construct tachometric curves conditioned on LIP activity, z-scored spike counts were divided into high and low spike count bins based on a median split. Tachometric curves were then constructed from trials within each bin. These analyses were performed on pooled data from the subpopulations of neurons analyzed in Fig. 5.

---

**Supplementary Figure 1.**
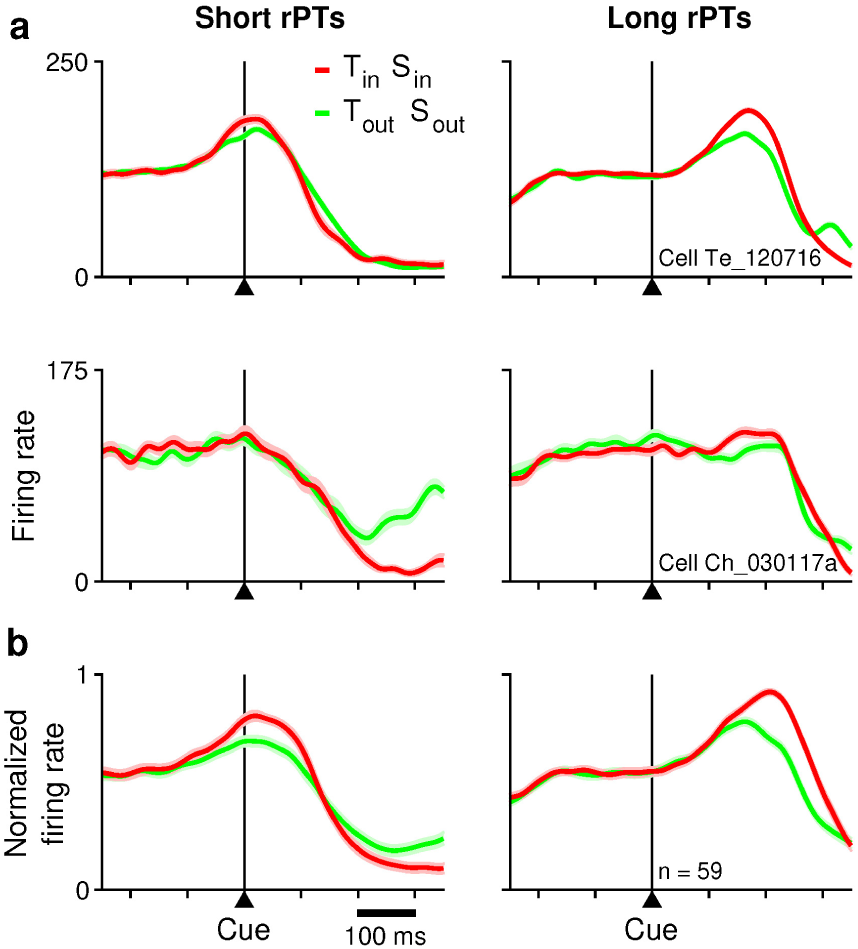
LIP neuronal discrimination aligned to cue onset. Same data and format as in Fig. 3, but with the recorded spikes aligned to cue onset.

**Supplementary Figure 2.**
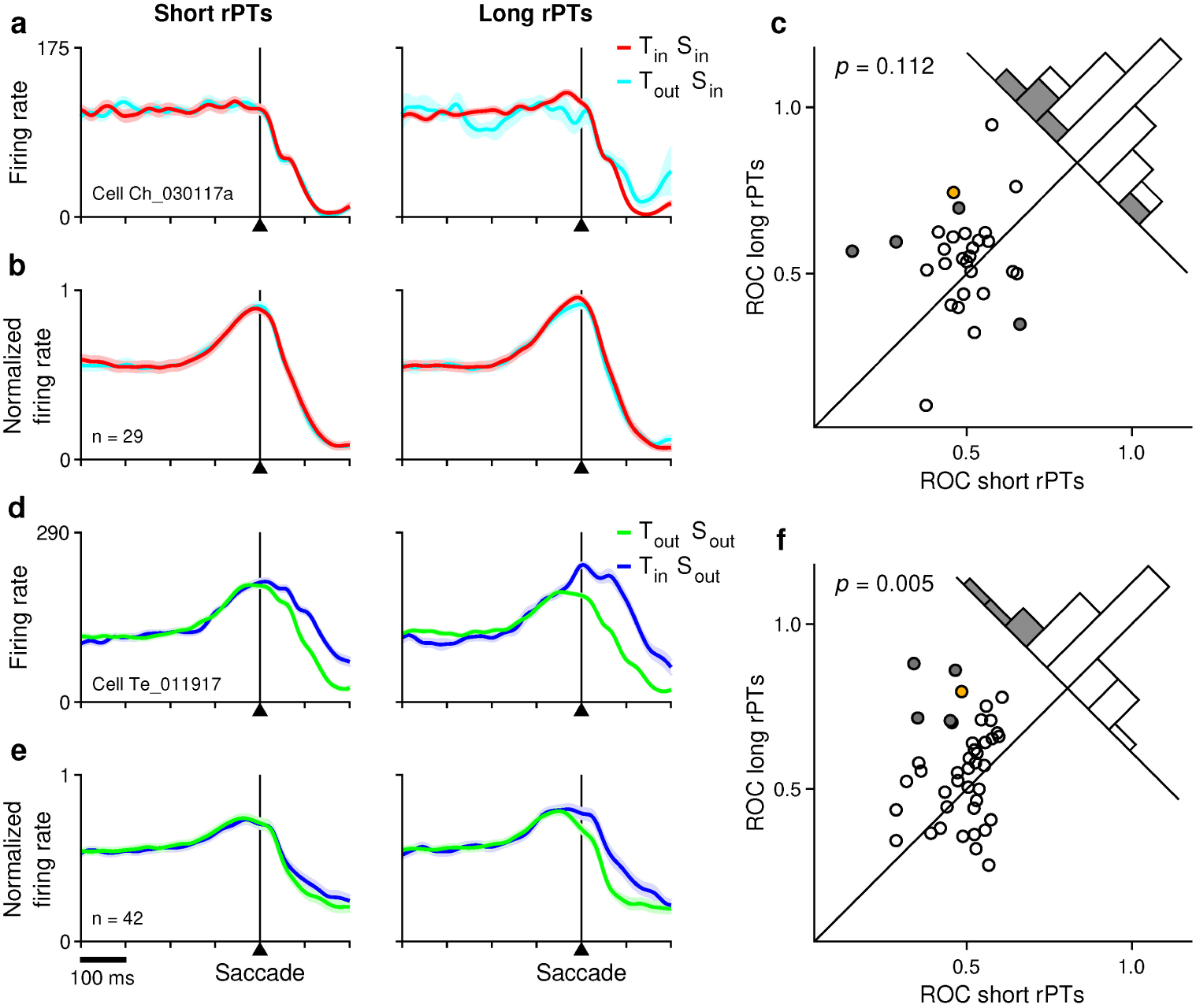
Cue-driven neuronal discrimination. Analyses are as in Fig. 3, but for choices with identical saccades. (**a**-**c**) Target-distracter discrimination during saccades into the RF. (**a**) Firing rate of a single LIP neuron as a function of time for correct T_in_ choices (target in, saccade in; red trace) and incorrect T_out_ choices (target out, saccade in; cyan trace) made at short (left) and long (right) rPTs. (**b**) Normalized population response from 29 LIP neurons, with identical conventions as in **a**. (**c**) ROC scores in long- (y-axis) versus short-rPT trials (x-axis). Each point represents data from one of the 29 neurons analyzed in this condition. Significance of median difference (Wilcoxon signed rank test) is indicated. Filled symbols and gray bars mark neurons with significantly different ROC scores (*p* < 0.05, permutation test). The colored point marks the example neuron in **a**. (**d**-**f**) Target-distracter discrimination during saccades away from the RF. Same format as in **a**-**c**, but for a group of 42 neurons analyzed in this condition.

